# Copper resistance in *Legionella pneumophila*: role of genetic factors and host cells

**DOI:** 10.1101/2024.09.04.611240

**Authors:** Gillian Cameron, Sebastien P. Faucher

## Abstract

Copper is frequently found in drinking water due to its presence in the natural environment and the widespread usage of copper pipes. This toxic metal has a well-known antimicrobial activity, an activity harnessed in copper-silver ionization (CSI) to eliminate the opportunistic pathogen *Legionella pneumophila* from engineered water systems. Despite utilizing the antimicrobial properties of copper in *Legionella* control, little is known about how copper containing environments affect *L. pneumophila* populations. The goal of this study is to understand how *L. pneumophila* responds to copper within a hot water distribution system (HWDS) environment. To answer this question, different sequence types and regulatory mutants were exposed to copper to compare their survival. *L. pneumophila* isolates of 4 sequence types from 3 different HWDSs exhibited a wide diversity of phenotypes after copper stress. The Δ*letA* and Δ*letS* mutants were sensitive to copper, indicating that the LetAS two component system is important for copper resistance. Additionaly, transmissive phase cultures were more resistant to copper than replicative phase cultures. Therefore, the regulation of entry into transmissive phase by the LetAS system is essential for *L. pneumophila’*s ability to survive copper stress. In a water system, *L. pneumophila* replicates within eukaryotic hosts. When cocultured with the host ciliate *Tetrahymena pyriformis*, *L. pneumophila* was more resistant to copper than when the bacteria were in a monoculture. No difference in *L. pneumophila* replication inside of hosts in cocultures with or without copper was observed. This result confirms that the presence of host cells protects *L. pneumophila* from copper stress. Therefore, presence of host cells in water system may limit the efficacy of copper-based control strategies.

## 1. Introduction

*Legionella pneumophila* is the causative agent of Legionnaires’ Disease, a severe form of pneumonia, and of Pontiac fever, a more flu-like illness (Cunha et al., 2016). Cases of Legionnaires’ Disease are increasing worldwide each year, making it a significant public health concern (Yu, Kamali, and Vugia, 2019). It is of particular concern as a nosocomial infection, as immunocompromised populations, the elderly, and people with comorbidities are at heightened risk of infection (Marston et al., 1994). In Canada, cases of legionellosis rose by 326% between 2010 and 2019, increasing from 140 cases per year to 655 cases per year (Government of Canada, 2021). However, the actual number of infections is likely much higher, as cases of Legionnaires’ Disease frequently go underreported, with an average of 2.8 illnesses, 2.5 hospitalizations, and 2.5 deaths for every reported Legionnaires’ Disease case, hospitalization, and death reported in Canada (McMullen et al., 2024). In the United States, a 2024 CDC report found *Legionella* infections were the leading cause of drinking water outbreaks and were responsible for 97% of hospitalizations and 98% of all deaths associated with waterborne pathogens between 2015 and 2020 (Kunz, 2024). *L. pneumophila* is found in freshwater environments and engineered water systems (EWSs), such as hot water distribution systems (HWDS) and cooling towers, where it is an obligate intracellular pathogen of protozoans, such as amoebas and ciliates (Fields et al., 1984; Rowbotham, 1980). *L. pneumophila* is transmitted to humans when water containing the bacteria is aerosolized from water distribution systems and inhaled (Cunha et al., 2016). Once inside of the lung, *L. pneumophila* is phagocytosed by alveolar macrophages, a process enhanced by the bacteria’s Icm/Dot Type IVb Secretion system to invade alveolar macrophages, causing infection and tissue damage (Escoll et al., 2014).

*L. pneumophila* has a biphasic life cycle, with an intracellular and extracellular phase (Byrne and Swanson, 1998; Molofsky and Swanson, 2004). The intracellular phase of *Legionella’*s life cycle is called replicative phase. During this phase the bacteria replicate to high levels within the host cell and are less resistant to stressors (Molofsky and Swanson, 2004). Once it has depleted its host of nutrients, *L. pneumophila* causes the lysis of the host cell and is released to the extracellular environment of a water system (Byrne and Swanson, 1998). This extracellular phase is known as the transmissive phase, during which *L. pneumophila* cells are more infectious and stress resistant, but do not replicate. The switch between replicative and transmissive phase is mediated primarily by the LetAS two component system (Hammer and Swanson, 2002). When *L. pneumophila* begins to deplete its host of nutrients and enter stationary phase, LetS, the sensor kinase, autophosphorylates (Rasis and Segal, 2009). It then phosphorylates the response regulator LetA which then activates the expression of RsmY and RsmZ, two non-coding small RNAs. RsmYZ bind to the post-transcriptional repressor CsrA (Rasis and Segal, 2009). Binding of CsrA by RsmY and RsmZ relieves its repression of the translation of mRNA, allowing transcription of genes involved in motility, virulence, and stress tolerance, including genes responsible for resistance to heat shock, oxidative stress, and acid stress (Molofsky and Swanson, 1999; Mendis et al., 2018).

Several treatment methods are used by water system operators to eliminate *L. pneumophila*. One such method is copper-silver ionization (CSI) (Liu et al., 1994). Both copper and silver are antimicrobial metals, and together have a synergistic effect (Lin, Stout, and Victor, 1996). For CSI, a copper anode and silver electrode are installed into a water system and periodically electrified to release metal ions into the water and eliminate any bacteria that may be present (LeChevallier, 2023). This treatment method is more effective at penetrating biofilms than chlorine and is less damaging to the water system as the metal ions do not cause corrosion. In addition to CSI, *L. pneumophila* can encounter copper from copper pipes, which are widely used in water distribution systems and have been found to select for chlorine-resistant bacteria (Khan et al., 2019). More generally, solid copper has been shown to select for copper resistant *Pseudomonas fluorescens* in an adaptive laboratory evolution study (Xu et al., 2022).

Copper has a well-characterized antimicrobial activity due to the high reactivity of Cu^+^ ions. When there is an excess of Cu^+^ ions inside of the cell, they will displace other metal ions from the core of metalloenzymes, rendering the enzymes non-functional (Giachino and Waldron, 2020). This mismetallation affects a wide variety of enzymes, disrupting cellular metabolic pathways and eventually leading to cell death. Copper ions also damage the cell membrane, increasing its permeability, though the precise mechanism of this damage remains unknown (Giachino and Waldron, 2020). Perhaps most famously, in aerobic conditions Cu^+^ ions react with oxygen to form toxic radical oxygen species (ROSs) (Solioz, 2018). These ROSs cause oxidative damage to the cell, disrupting redox potential and damaging the DNA. Despite initial success in using CSI in their HWDS to reduce the number of Legionnaires’ Disease cases to zero, some users have reported the reemergence of Legionnaires’ Disease (Demirjian et al., 2015).

Investigation showed that *L. pneumophila* isolated from these water distribution systems remained viable when exposed to both copper and silver (Demirjian et al., 2015).

Currently, the only characterized copper resistance gene in *L. pneumophila* is *copA* (Kim et al., 2009; Trigui et al., 2013). CopA is a P-type ATPase pump, coupling the transport of Cu^+^ ions from the cytoplasm to the periplasm with ATP hydrolysis (Kim et al., 2009). Inside of the periplasm, Cu^+^ ions are converted into more stable Cu^2+^ ions by a multicopper oxidase, such as *Escherichia coli*’s CueO. Multiple putative multicopper oxidases have been identified in *L. pneumophila* such as *mcoL* (Huston et al., 2008). CopA is an example of the acquisition of resistance genes by pathogens through horizontal gene transfer, as *copA* is encoded on the pLP100 mobile genetic element (Trigui et al., 2013). Copper resistance has also been observed to emerge through adaptation to a copper-containing microenvironment via the accumulation of point mutations. Bédard et al. isolated four isolates of *L. pneumophila* strain ST1427 from HWDS in a building with an ongoing Legionnaires’ Disease outbreak (Bédard et al., 2021). Two of the isolates were sensitive to copper, while the other two, from a biofilm in a copper pipe within the same system, were tolerant to copper. The tolerant and sensitive isolates differed by only 29 single nucleotide polymorphisms (SNPs), indicating that the *L. pneumophila* isolates had adapted to their local microenvironment through the accumulation of point mutations. Of the 29 SNPs identified, only 1 occurred near a known copper resistance gene (*copA)*, suggesting the presence of genes with a previously unknown role in *L. pneumophila*’s response to copper stress.

The aim of this study is to deepen our understanding of how *L. pneumophila* survives copper stress. We examined environmental and clinical isolates from different HWDSs to determine the phenotypic diversity of resistance to copper stress in *L. pneumophila.* Then we examined the genes that may be involved in copper resistance, hypothesizing that *L. pneumophila*’s general stress response played a role in copper resistance. And finally, we investigated the ability of host cells to protect *L. pneumophila* from the toxic effects of copper.

## 2. Materials and Methods

### 2.1 *Culturing* L. pneumophila

*L. pneumophila* strains (Table 1) were kept as frozen stocks at −80⁰C. Strains were grown on CYE (ACES-buffered charcoal yeast extract) agar plates adjusted to pH 6.9 with 10 mM KOH and containing 0.25g/ L-cysteine and 0.4g/L ferric pyrophosphate supplements at 37⁰C for 3 days (Feeley et al., 1979). When needed, media was supplemented with 5 mg/mL chloramphenicol, 25 mg/mL kanamycin sulfate, 15 µg/mL gentamicin, or 0.1 mM isopropyl β-d-1- thiogalactopyranoside (IPTG). Individual colonies were grown in 1 mL of AYE broth (CYE lacking charcoal or agar) at 37⁰C with shaking. IPTG was added to cultures and suspensions of SPF39 and SPF312, at a final concentration of 0.1 mM to induce expression of LetA and LetS, respectively.

**Table 1:** Average slope and standard deviation of transmissive and replicative phase suspensions of four biological replicates after exposure to CuCl2.

|  | Transmissive<br>0 mM | Replicative<br>0 mM | Transmissive<br>0.08 mM | Replicative<br>0.08 mM | Transmissive<br>0.16 mM | Replicative<br>0.16 mM |
| --- | --- | --- | --- | --- | --- | --- |
| Slope | -0.05657 | -0.02242 | -0.2572 | -0.4114 | -0.4246 | -0.6109 |
| Standard<br>Deviation | 0.009482 | 0.01331 | 0.026 | 0.03583 | 0.03587 | 0.05994 |

### 2.2 Copper exposure and survival tests

Overnight cultures were pelleted at 5000×*g* for 5 minutes, then washed three times in Fraquil, a defined low-nutrient medium that simulates freshwater (Morel at al., 1975) and suspended in Fraquil. Cell concentration was determined by measuring the OD600 with a spectrophotometer. Bacterial suspensions were diluted to a final OD600 of 0.1, equivalent to about 1x10^8^ cells/mL. The diluted suspensions were incubated overnight at room temperature. In a 24 well plate, 10 µL of 8 mM CuCl2 was added to each well for a final concentration of 0.08 mM CuCl2. For the control conditions, 10 µL of Fraquil was added to each well. 990 µL of the bacterial suspensions were added to each well and left at room temperature for 4 hours. After the 4-hour incubation period, bacterial cultures were diluted and plated onto CYE. A CFU count was used to determine survival.

### 2.3 Comparison of transmissive and replicative phase

For a culture of E phase cells, a Philadelphia-1 culture was grown in 1 mL AYE media and shaken at 37⁰C for 12 hours. To generate a PE phase culture of *L. pneumophila,* and overnight culture of the *L. pneumophila* strain Philadelphia-1 was grown in 1 mL AYE media and shaken at 37C for 48 hours. Both cultures were pelleted at 8,000xg for 3 minutes and washed and resuspended in 2 mL Fraquil 3 times. Bacterial suspensions were diluted to an OD600 of 0.1, equivalent to 1x10^8^ cells/mL, then left at room temperature overnight. 10 µL of Fraquil, 8mM CuCl2, or 16 mM CuCl2 were added to wells in a 24 well plate. 990 µL of transmissive phase suspension or replicative phase suspension were added to each well. CFU counts of the bacterial suspensions with copper were taken at 0, 1, 4, and 8 hours to determine bacterial reduction over time.

### 2.4 Host cell maintenance and coculture

*T. pyriformis* was stored at −80⁰C. To start cell cultures from these frozen stocks, stocks were thawed at 37⁰C with SPP+ 8% sucrose (proteose peptone, dextrose, yeast extract, FeCl, water, and sucrose), then grown in PYNFH+ 4% sucrose+10% FBS overnight (ATCC medium 1034 with sucrose and heat inactivated FBS). Cultures were then transferred to ATCC 357 media (protease peptone, tryptone, and potassium phosphate) for maintenance. 3 mL *T. pyriformis* cultures in 20 mL ATCC 257 media were passaged weekly. Prior to infection, 3 mL of *T. pyriformis* culture was passaged in 20 mL of complete modified PYNFH media (ATCC medium 1034) with FBS for 3 days. For infection, *T. pyriformis* cultures were pelleted at 600×*g* for 5 minutes, then washed with Fraquil. A sample of the cells were stained with 0.4% Trypsin blue and counted with a hemocytometer. The washed *T. pyriformis* cells were diluted to 5x10^5^ cells/mL in Fraquil. *L. pneumophila* strains were resuspended in Fraquil and diluted to an OD600 of 0.1 (1x10^8^ cells/mL). To set up cocultures, 1 mL of diluted *T. pyriformis* cells was added to each well in a 24 well plate. 5 µL of *L. pneumophila* suspensions were added to each well, for a final concentration of 1x10^5^ cells/mL. For the copper exposed condition, 10 uL of 8 mM CuCl2 was added. For the control condition, the cocultures were diluted with 10 uL of Fraquil. CFU counts of each infection were taken every 24 hours. A coculture of *T. pyriformis* and the *L. pneumophila* ΔdotA mutant was used as an infection control.

## 3. Results

### 3.1 Resistance of L. pneumophila isolates to copper

A variety of clinical and environmental isolates of *L. pneumophila* of different sequence types (ST) from 3 different HWDSs were tested for survival in the artificial freshwater Fraquil in the presence of copper (Fig. 1; Morel et al., 1975). The isolates were chosen because of their previous exposure to high levels of copper. Except for SPF635, all ST378 isolates were taken from a HWDS in site A, a healthcare facility built in 1971 (Table 2; Bédard et al., 2019). This HWDS utilizes copper pipes for all plumbing (Bédard et al, 2015). Because of the copper pipes, site A’s HWDS had high levels of copper, with an average of 478 µg/L and a maximum of 743 µg/L (Bédard et al., 2019). Site B, where SPF635 was isolated from, is a healthcare facility built in 1903. Site C, where the ST2859 isolates were taken, is a recreational centre built in 1976. The HWDS of site C also utilized copper pipes and had high levels of copper, with an average of 2764 µg/L. Because these sites had high concentrations of copper in the HWDS, we wanted to determine if this previous exposure to copper selected for copper resistance in these strains compared to the lab strain *L. pneumophila* Philadelphia-1. A bioinformatics analysis of the ST378 isolates showed that these isolates have 2 copies of the P-type ATPase copper efflux pump *copA* on their chromosome, while *Philadelphia-1* only has1 copy (Najeeb et al., 2024, submitted). All isolates tested were significantly affected by CuCl2 exposure but were not significantly different from the wildtype (Fig. 1A). Only two isolates, SPF546 and SPF547 were significantly different from the Philadelphia-1 type strain (P=0.0036 and P=0.0042, respectively) (Fig. 1B). Overall, the survivability to copper seems to be variable amongst the strains tested, with no clear correlation with the source of isolation and the concentration of copper in the water system.

**Figure 1:**
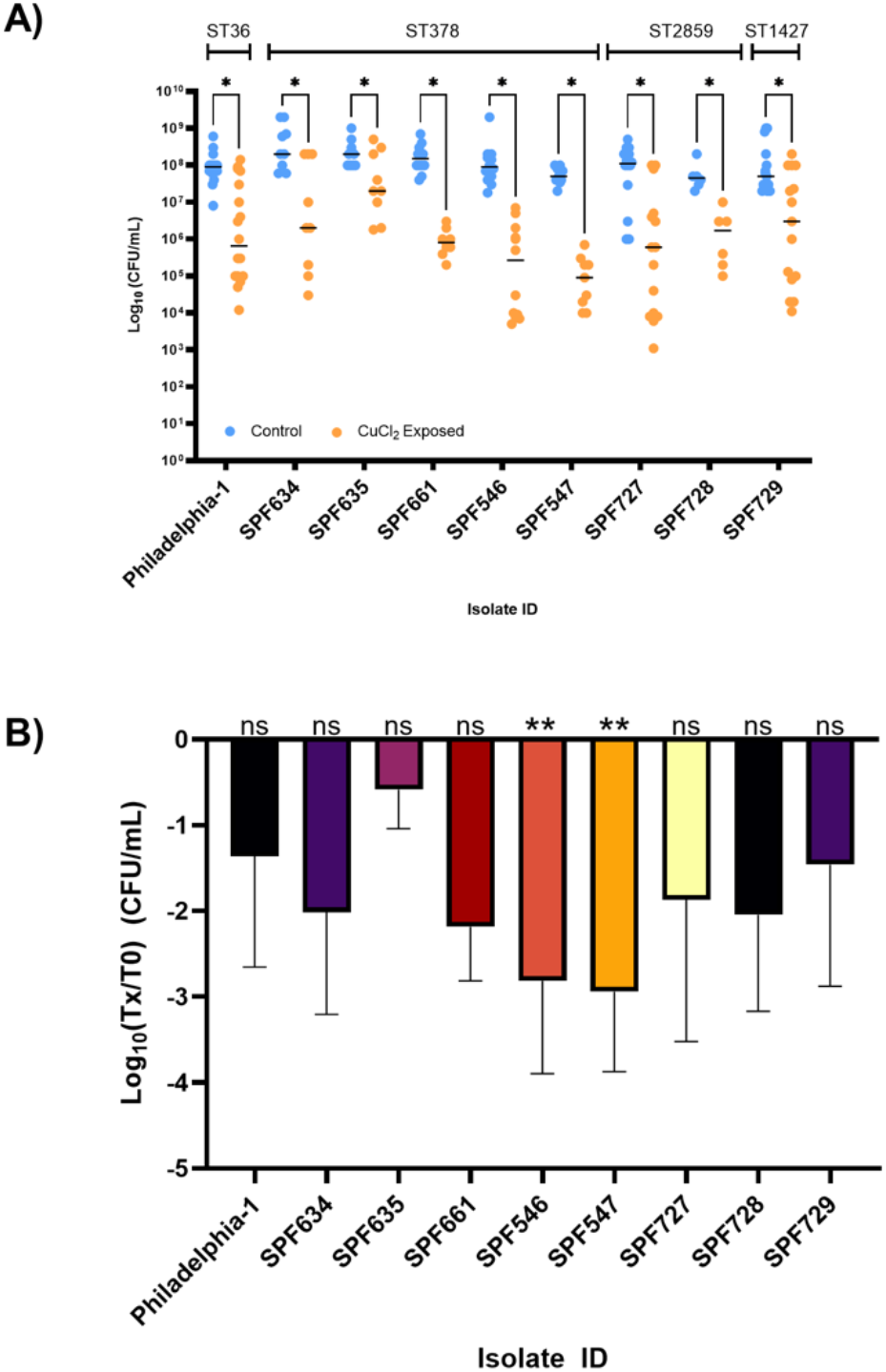
Effect of copper on environmental *L. pneumophila* isolates. Strains were grown in overnight broth culture, then washed and resuspended in Fraquil to a final concentration of 10^8^ cells/mL. Suspensions were incubated overnight at room temperature, then exposed to 0.08 mM CuCl2. Control suspensions were not exposed to copper. After incubation at room temperature for 4 hours, survival was determined using viable cell counts on CYE agar medium. A) Raw CFU count data. Significant effect of copper on survival of each strain was determined using paired, non-parametric t tests, *P* <0.05 indicated with *, ns represents a non-significant result. Data shown are the values of individual replicates with the black line indicating the average. B) Ratio of CuCl2:Control CFU counts. Data was analyzed with an uncorrected ordinary one-way ANOVA, then analyzed with multiple comparisons. Significant comparisons (*P* <0.05) indicated with an * and non-significant comparisons indicated with ns.

**Table 2:** *L. pneumophila* strains and isolates used in this study.

| Isolates ID | Description | SBT | Source<sup>1</sup> | Isolation Date | Ancestor |
| --- | --- | --- | --- | --- | --- |
| Philadelphia-1 |  | ST36 | ATCC 33152 | - | - |
| KS79 | JR32 $\Delta comR$ | - | (de Felipe et al., 2008) | - | JR32 |
| SPF186 | $\Delta copA:GT^R$ | - | (Kim et al., 2009) | - | KS79 |
| SPF605 | lpg0563(T1) $phaP \Delta Int39 :Kn^R$ | - | (Liang, Cameron, and Faucher, 2023) | - | KS79 |
| SPF159 | $\Delta cpxR:Kn^R$ | - | (Tanner et al., 2016) | - | KS79 |
| $\Delta letS$ | $\Delta letS: Kn^R$ | - | (Hovel-Miner et al., 2009) | - | KS79 |
| SPF39 | $\Delta letS+pSF21:CM^R$ | - | (Hovel-Miner et al., 2009) | - | $\Delta letS$ |
| $\Delta letA$ | $\Delta letA:Kn^R$ | - | (Gal-Mor and Segal, 2003) | - | KS79 |
| SPF312 | $\Delta letA+pMMB207-letA:CM^R$ | - | (Lynch et al., 2003) | - | $\Delta letA$ |
| SPF544 | Environment | ST378 | A (Najeeb et al., 2024, submitted) | 2021 | - |
| SPF546 | Environment | ST2859 | C (Matthews et al., 2022) | 2021 | - |
| SPF547 | Environment | ST2859 | C (Mathhews et al., 2022) | 2021 | - |
| SPF597 | Environmental<br>(Taken after 1 round of heat disinfection) | ST378 | A (Najeeb et al., 2024, submitted) | 2021 | - |
| SPF599 | Environment | ST378 | A (Najeeb et al., 2024, submitted) | 2021 | - |
| SPF600 | Environment | ST378 | A (Najeeb et al., 2024, submitted) | 2021 | - |
| SPF601 | Environment | ST378 | A (Najeeb et al., 2024, submitted) | 2021 | - |
| SPF634 | Environment | Similar to ST378 | A (Najeeb et al., 2024, submitted) | 2015 | - |
| SPF635 | Environment (Hot water tap) | Similar to ST378 | B (Najeeb et al., 2024, submitted) | 2015 | - |
| SPF652 | Environment | ST378 | A (Najeeb et al., 2024, submitted) | 2012 | - |
| SPF661 | Environment | ST378 | A (Najeeb et al., 2024, submitted) | 2013 | - |
| SPF727 | Environment | ST2859 | C (Matthews et al., 2022) | 2020 | - |
| SPF728 | Environment | ST2859 | C (Matthews et al., 2022) | 2020 | - |
| SPF729 | Environment | ST1427 | C (Matthews et al., 2022) | 2020 | - |

1) For isolates SPF544-SPF729, the site of isolation is indicated by letters: Site A, B and C.

Sites A, B, and C all had high concentrations of copper within their HWDSs. When environmental isolates of *L. pneumophila* taken from these systems were exposed to copper, a large diversity in the isolates’ ability to survive copper stress was observed (Fig. 1A). A variety of responses to copper stress was observed in the ST2859 isolates, with SPF546 and SPF547 less resistant to copper than the other isolates. In general, there appears to be a wide diversity in susceptibility to copper stress among *L. pneumophila* strains, even among different isolates of the same sequence type taken from the same HWDS. This diversity could be attributable to differences in microenvironments within the water system. A HWDS is not uniform environment; there are dead end pipes, differing flow rates, and different water compositions depending on pipe material. Bédard et al. (2021) found differences in pipe microenvironment to play a large role in copper resistance in *L. pneumophila* (Bédard et al., 2021). Isolates taken from a biofilm in a copper pipe were copper resistant, while isolates from the water were sensitive.

The resistant isolates had adapted to their copper-rich microenvironment within the biofilm. Because they lived in an environment that had a lower concentration of copper, the water isolates had not adapted to survive copper stress. It is possible that within site A and within site B, differences in flow rate or the presence of dead-end pipes created microenvironments that had locally higher or lower concentrations of copper, resulting in isolates of *L. pneumophila* with a wide diversity in survival in the presence of copper.

### 3.2 CpxRA is unnecessary for copper resistance in L. pneumophila

To understand what genes may be involved in adaptation to a copper-containing microenvironment, the survival of *L. pneumophila* mutants lacking stress regulators in the presence of copper was tested. It was expected that a *ΔcpxR* mutant would be significantly more susceptible to copper than the wild type, as in other Gram-negative species, such as *E. coli,* the CpxRA two component system responds to copper stress, specifically to the damage to the cell envelope caused by copper ions (Yamamoto and Ishihama, 2006). One of the proteins regulated by CpxRA is the tail specific protease (Tsp) (Saoud, Mani, and Faucher, 2021). In other bacteria, Tsp modulates peptidoglycan synthesis, and in *L. pneumophila* has been shown to be important for surviving thermal stress (Lawrence et al., 2014; Singh et al., 2015; Saoud, Mani, and Faucher, 2021). We hypothesized that misregulation of membrane homeostasis affects copper sensitivity in *L. pneumophila*. The susceptibility to copper of Δ*cpxR*, a mutant in the CpxRA two component system responsible for the envelope stress response, and Δ*tsp*, which lacks *L. pneumophila*’s tail specific protein, was tested. In other Gram-negative species, the CpxRA two component system responds to envelope damage induced by copper. As such, we expected that the ΔcpxR mutant would show increased sensitivity to copper. However, survival of the ΔcpxR mutant was not significantly different from the WT, indicating that the CpxRA system has no role in copper resistance in *L. pneumophila* at least in the conditions tested (Fig. 2). Since copper ions also cause the misfolding of proteins, the ability of a Δtsp mutant to survive in copper was tested. Like Δ*cpxR*, the Δ*tsp* mutant showed no significant difference in survival after exposure to copper from the WT, indicating Tsp does not play a role in *L. pneumophila*’s response to copper stress (Fig. 2).

**Figure 2:**
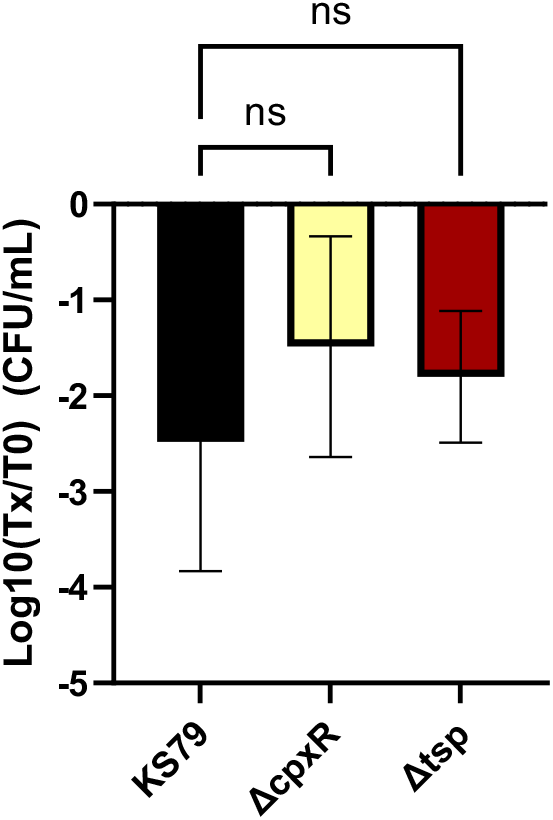
Effect of copper on *L. pneumophila* mutants in membrane stress response regulators. Strains were grown in overnight broth culture, then washed and resuspended in Fraquil to a final concentration of 10^8^ cells/mL. Suspensions were incubated overnight at room temperature, then exposed to 0.08 mM CuCl2. Control suspensions were not exposed to copper. After incubation at room temperature for 4 hours, survival determined using viable cell counts on CYE agar medium. Data shown represent average and standard deviation of 3 biological replicates. Statistically significant difference from the WT (KS79) determined with one-way ANOVA and multiple comparisons, *\** indicates *P* < 0.05.

### 3.3 LetA/S two component system has a role in copper resistance

The biphasic lifestyle of *L. pneumophila* is important for stress resistance (Bachman and Swanson, 2001; Lynch et al., 2003). The effect of mutation in *letA* and *letS* on copper resistance was therefore investigated. When analyzed with one-way ANOVA and Tukey’s multiple comparisons, survival of *ΔletA* and *ΔletS* were significantly different (*P*<0.0001) from the wildtype KS79 (Fig. 3A). Complementation study was then carried out to confirm the role of these genes. The susceptibility of the merodiploid strains, SPF39 and SPF312, carrying a *letS* and a *letA* genes driven by the *Ptac* promoter, respectively, were tested. When uninduced, SPF312 was a bit more resistant to copper than the Δ*letA* mutant (*P* <0.0001), but SPF39 was no different than the Δ*letS* mutant (*P =* 0.044) (Fig. 3A). When induced with 0.01 mM IPTG and compared with Tukey’s multiple comparisons, both SPF39 and SPF312 were significantly more resistant than the corresponding Δ*letS* and Δ*letA* mutant strains (*P*=0.0014 and *P*<0.0001, respectively) and return to wild-type levels (Fig. 2C). These results indicate that the LetAS two- component system plays a significant role in copper resistance in *L. pneumophila*.

**Figure 3:**
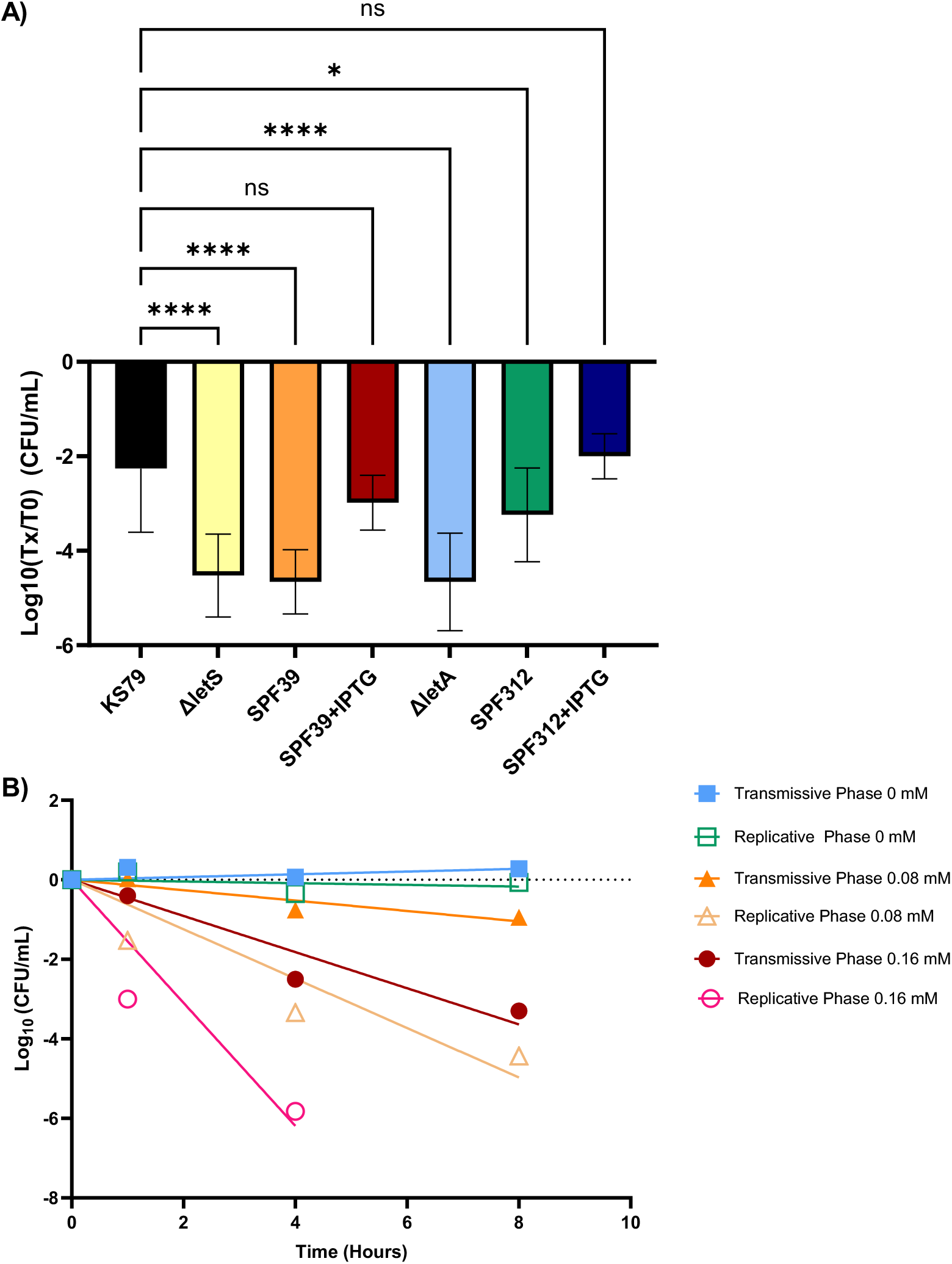
The LetAS regulator is required for copper resistance. A) Survival ratios of *L. pneumophila* after exposure to 8 mM CuCl2 for 4 hours. Strains were grown in overnight broth culture, then washed and resuspended in Fraquil to a final concentration of 10^8^ cells/mL. Suspensions were incubated overnight at room temperature, then exposed to 0.08 mM CuCl2. Control suspensions were not exposed to copper. After incubation at room temperature for 4 hours, survival determined using viable cell counts on CYE agar medium. Expression of LetS and LetA in SPF39 (ΔletS + pletS) and SPF312 (ΔletA + pletA), respectively, was induced with 0.1 mM IPTG. Data shown represent the average and standard deviation of 3 biological replicates. Statistically significant difference from the WT (KS79) was determined with one-way ANOVA and multiple comparisons, *\** indicates *P* < 0.05. B) Reduction in cell concentration over time of *L. pneumophila* Philadelphia-1 in transmissive phase and replicative phase. The transmissive phase culture was grown in a shaker at 37⁰C for 48 hours, while the replicative phase culture was grown in a shaker 37⁰C for 12 hours to reach replicative phase. Both cultures were washed and resuspended in Fraquil to a final concentration of 10^8^ cells/mL and incubated at room temperature for 24 hours, then exposed to 0 mM, 0.08 mM, and 0.16 mM CuCl2. CFU counts were taken at 0, 1, 4, and 8 hours. Ration against the T0 was calculated, transformed, and fitted with linear regression. Figure shows one representative replicate. See supplemental data for individual replicate data.

Two possible options could explain the increased sensitivity of the Δ*letA* and Δ*letS* mutants to the effect of CuCl2: A) the mutants were unable to activate expression of copper resistance genes, or B) as the LetAS system regulates entry into the transmissive phase of *L. pneumophila’* s life cycle, the Δ*letA* and Δ*letS* strains were unable to enter transmissive phase, which is more stress tolerant, and were therefore more sensitive to the effect of copper. A previous study by Mendis et al. (2018) examined the genes activated by LetA, but none of these genes were known to have a role in copper resistance (Mendis et al., 2018). The increased copper sensitivity in the Δ*letA* and Δ*letS* mutants is likely due to an inability to switch from replicative phase to transmissive phase. Therefore, the effect of copper on transmissive phase and a replicative phase culture of *L. pneumophila* was compared. *L. pneumophila* Philadelphia-1 cultures in transmissive phase and replicative phase were exposed to 0, 0.08 and 0.16 mM CuCl2. CFU counts were taken at 0, 1, 4, and 8 hours to compare survival. The ratio against the baseline values of 0 mM CuCl2 was calculated, transformed, and a line of best fit was graphed. At 0.08 mM CuCl2, the transmissive phase suspension had a slope of −0.2572, while the replicative phase suspension had a slope of −0.4114 (Fig. 3B; Table 1; Supplementary Figure 1). When exposed to 0.16 mM CuCl2, the transmissive phase suspension had a slope of −0.4246, while the replicative phase suspension had a slope of −0.6109 (Fig. 3B; Table 1; Supplementary Figure 1). At all concentrations tested, the slope of the transmissive phase suspension was steeper than the slope of the replicative phase suspension, indicating the transmissive phase suspension survived better in the presence of copper (Fig. 3B; Table 1; Supplementary Figure 1). This confirms *L. pneumophila* is more resistant to copper when it is in transmissive than when it is in replicative phase. Taken together, these results indicated that LetAS regulation of entry into transmissive phase is important for copper resistance in *L. pneumophila*.

The results of this study are consistent with Lynch et al. (2003), who found that *letA* mutants were defective in the stationary phase stress response, and were sensitive to oxidative stress (Lynch et al., 2003). As copper reacts with oxygen to form toxic oxygen radicals, it is possible the Δ*letA* and Δ*letS* mutants are more sensitive to the oxidative stress caused by copper phase (Lynch et al., 2003; Hammer, Tateda, and Swanson, 2002). This would also explain the increased sensitivity of *L. pneumophila* in replicative phase to copper stress compared to *L. pneumophila* in transmissive phase. The LetAS system does not directly regulate known copper resistance genes in *L. pneumophila* (Mendis et al., 2018). However, LetAS does regulate the expression of genes involved in oxidative stress resistance, namely *sodB*, which encodes a superoxide dismutase, and *ohr*, which encodes an organic hydroperoxide resistance protein (Mendis et al., 2018). Both of these genes are involved in dismantling oxygen radicals to protect *L. pneumophila* from oxidative stress (Broxton and Culotta, 2016; Brown, 2019). When the ability of transmissive phase and replicative phase cultures to survive in the presence of copper was compared, replicative phase cultures were more sensitive to copper stress than transmissive phase cultures at both concentrations tested (Fig. 3B). Based on this result, the most likely explanation for the sensitivity towards copper stress observed in the *ΔletA* and *ΔletS* mutants is that without a functioning LetAS system, the mutant cells were stuck in replicative phase and unable to enter the more stress tolerant transmissive phase, increasing their sensitivity to copper. This result is consistent with the findings of Sahr et al. (2017). In their study, Sahr et al found that the Δ*letA* mutant had a defect in entering an *Acanthamoeba castellanii* cell, but was unable to replicate inside of the host, indicating that without LetA the mutant was halted in replicative phase (Sahr et al., 2017). The increased copper sensitivity of the *ΔletA* and *ΔletS* mutants is also consistent with the findings of Hammer, Tateda, and Swanson (2002), who found that *letA* and *letS* mutants were unable to express transmissive phase traits (Hammer, Tateda, and Swanson, 2002). These mutants were non-motile, non-cytotoxic, and had poor efficiency for the infection of macrophages (Hammer, Tateda, and Swanson, 2002) The LetAS system influences copper sensitivity, possibly via control of oxidative stress resistance genes and the activation of other general stress responses.

### 3.4 Presence of host cells protect L. pneumophila from the effects of copper

Next, we investigated the effect of intracellular growth on the susceptibility to copper. Cocultures of *L. pneumophila* and the ciliate species *Tetrahymena pyriformis* were set up in Fraquil with an MOI of 1 (Fig. 4). Control cocultures received 10 µL of Fraquil while treated coculture received 0.08 mM CuCl2. In parallel, the survival of *L. pneumophila* without host cells was also tested. A CFU count was taken every 24 hours to quantify *L. pneumophila* replication, and the CFU counts were compared to the WT using an unpaired, one-tailed student’s t-test.

**Figure 4:**
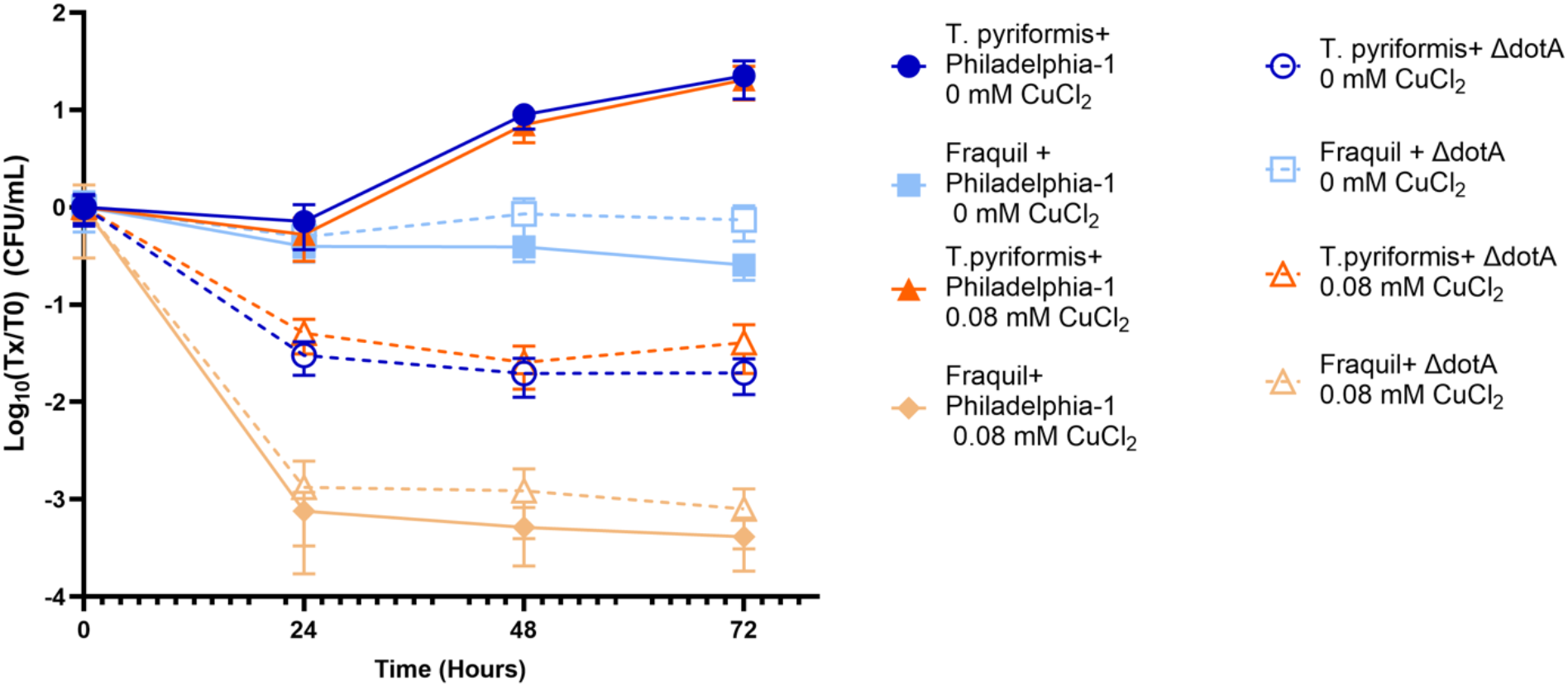
Effect of copper on *L. pneumophila* in host cells. 10 uL of 8 mM CuCl2 was added directly to cocultures of *L. pneumophila* and *T. pyriformis.* The *L. pneumophila* strain ΔdotA was used as an infection control. Growth of *L. pneumophila* strains Philadelphia-1 and ΔdotA in the presence of copper in Fraquil and in coculture with *T. pyriformis*. MOI of 1. Samples were taken every 24 hours and plated on CYE agar plates for a CFU count to determine cell survival over time. ΔdotA control for each condition represented with a dotted line. Data represents average and standard deviation of 3 biological replicates. Data was analyzed with an unpaired, one-tailed Student’s t-test to assess statistical significance versus the WT.

Without host cells, the number of cells of *L. pneumophila* strains Philadelphia-1 and *ΔdotA* rapidly decreased in the presence of copper (Fig. 4, light orange solid dotted lines), but stayed relatively constant without copper (light blue solid and dotted lines), as expected. When cocultured with *T. pyriformis* Philadelphia-1 showed good replication in the presence of 0.08 mM CuCl2, similar to growth observed in coculture without CuCl2. Replication inside of host cells in the presence of 0.08 mM CuCl2 was not significantly different from replication without CuCl2 (P=0.4423) when analyzed with an unpaired, one-tailed t-test. When cocultured with *T. pyriformis, ΔdotA* showed a slight decrease over time in both the copper exposed and the control condition (Fig. 4, compare dark orange dotted line with dark blue dotted line); but survive much better that without host cells (Fig. 4, orange dotted line). Taken together, our results show that *L. pneumophila* can efficiently grow within *T. pyriformis* in the presence of copper and that the presence of host cell somewhat protects *L. pneumophila*.

Our examination of the ability of different *L. pneumophila* isolates and mutants to survive in the presence of copper had primarily focused on bacterial cells surviving independently in water. But in the HWDS environment, *L. pneumophila* does not replicate freely in the water, instead replicating within protozoan hosts (Kwaik et al., 1998; Rowbotham; 1980). Initially, both the amoeba *V. vermiformis* and *T. pyriformis* were used for this experiment and PYNF media was used in the cocultures. Both host cell types survived well in this media, but the PYNF media attenuated copper toxicity (Supplementary Figure S2). Presumably, the copper ions bound to components of the media due to their high reactivity. Previous coculture experiments with *T. pyriformis* and *L. pneumophila* used sterile tap water, so we selected Fraquil as the media for cocultures to avoid this issue (Fields et al., 1984). However, in Fraquil *V. vermiformis* cells died before *L. pneumophila* could infect them. Only *T. pyriformis* was used for this experiment as we could not find a system compatible with both *V. vermiformis* and copper.

There was no significant difference in *L. pneumophila* replication within a ciliate host between cocultures of *T. pyriformis* and *L. pneumophila* Philadelphia-1 with and without CuCl2, while survival of both Philadelphia-1 and *ΔdotA* in single species suspensions with copper decreased significantly within 24 hours, indicating that the presence of host cells protects the bacteria from copper stress. The role of host cells in protecting *L. pneumophila* from harmful environmental stressors has been previously observed. Storey et al. (2004) demonstrated that interaction between *Acanthamoebae* hosts and Legionella protects the bacteria from thermal treatment of water systems (Storey et al., 2004). However, interactions between amoeba hosts and *L. pneumophila* was shown to increase sensitivity to chlorination. Unlike amoebas, however, ciliates like *T. pyriformis* do not form protective cysts to survive environmental stresses (Salazar-Ardiles, Asserella-Rebollo, and Andrade, 2022; Nanney and McCoy, 1976; Lynn and Doerder, 2012). It is likely that the host cells are buffering the available copper. Copper ions are highly reactive, and readily bind to a wide variety of substrates (Solioz, 2018). It is likely the Cu^2+^ ions added into the coculture in the form of CuCl2 bound to the negatively charged cell membrane of *T. pyriformis*, reducing the amount of available copper. This would explain why the Δ*dotA* mutant was able to persist at low levels in coculture with *T. pyriformis* and CuCl2 but dies off when in a monoculture with copper. The possible ability of ciliates to attenuate available copper ions will need to be confirmed in a future study. Compared to *L. pneumophila, T. pyriformis* are less sensitive to CuCl2: concentrations higher than. 200 mg/L (equivalent to 1.48 mM) are required to inhibited *T. pyriformis* growth (Niclau et al., 1999). This is much higher than the concentration of CuCl2 needed to inhibit *L. pneumophila,* which has an MIC of around 0.5 mM (Supplementary Figure 3). Because of the difference in concentration of copper required to kill the bacteria and *T. pyriformis,* it is also possible *T. pyriformis* shelters *L. pneumophila* from copper toxicity. At CuCl2 concentrations that are subinhibitory for ciliates, *L. pneumophila* cells inside of the ciliate would be protected from the effects of copper toxicity.

Protection of *L. pneumophila* by hosts from copper toxicity could also explain the diversity in responses to copper stress observed in the ST378 and ST2859 isolates (Fig. 1). Different microenvironments within the HWDSs could favour the growth of host cells, such as dead-end pipes, which have lower flow rates than the rest of the system. The higher concentration of hosts within these microenvironments could protect *L. pneumophila* from copper toxicity, thus removing the selective pressure applied on the bacteria by copper. As such, *L. pneumophila* isolates from these environments would be less resistant to copper than isolates from microenvironments with less host cells to protect the bacteria. Niclau et al. (1999) found that subinhibitory concentrations of copper stimulate grazing by *T. pyriformis* (Niclau et al., 1999). The presence of copper within different microenvironments within the built environment would increase grazing by ciliate hosts, increasing protection of *L. pneumophila* as uptake of bacteria increased. This could also explain why environments with high concentration of copper still harbor *L. pneumophila*.

## 4. Conclusions

Copper in freshwater environments within EWSs has a major impact on *L. pneumophila*, whether that be copper leaching from copper pipes or copper ions released in CSI to control bacterial populations. When exposed to 0.08 mM CuCl2 environmental isolates from three different copper-containing HWDSs had diverse copper resistance phenotypes. No correlation between location, isolate source, or sequence type and copper resistance could be determined. When mutants lacking stress regulators were exposed to copper, it was found that the LetAS system is essential for copper resistance. This is likely because entry into the transmissive phase of *L. pneumophila*’s biphasic life cycle is required for copper resistance, as cultures of *L. pneumophila* in transmissive phase were more resistant to copper than cultures in transmissive phase. When *L. pneumophila* was cocultured with *T.pyriformis,* the presence of the host cell sheltered the bacteria from the stressor, as there was no significant difference between cocultures with and without copper.

## Acknowledgements

This study was funded by the Natural Sciences and Engineering Research Council of Canada (NSERC) Discovery grant RGPIN-2024-04228 to S.P.F. G.C. acknowledges the support of the NSERC Collaborative Research and Training Experience (CREATE) One Health Against Pathogens program.

## Bibliography

1. Abu Kwaik, Y., Gao, L. Y., Stone, B. J., Venkataraman, C., & Harb, O. S. (1998). Invasion of protozoa by Legionella pneumophila and its role in bacterial ecology and pathogenesis. Applied and environmental microbiology, 64(9), 3127–3133.

2. Bédard, E., Fey, S., Charron, D., Lalancette, C., Cantin, P., Dolcé, P., … & Prévost, M. (2015). Temperature diagnostic to identify high risk areas and optimize Legionella pneumophila surveillance in hot water distribution systems. Water research, 71, 244–256.

3. Bédard, E., Paranjape, K., Lalancette, C., Villion, M., Quach, C., Laferrière, C., … & Prévost, M. (2019). Legionella pneumophila levels and sequence-type distribution in hospital hot water samples from faucets to connecting pipes. Water research, 156, 277–286.

4. Bédard, E., Trigui, H., Liang, J., Doberva, M., Paranjape, K., Lalancette, C., … & Prévost, M. (2021). Local adaptation of Legionella pneumophila within a hospital hot water system increases tolerance to copper. Applied and environmental microbiology, 87(10), e00242–21.

5. Brown, C. L. (2019). Secretion and Environmental Biochemistry of Legionella pneumophila in Corrosive Water (Doctoral dissertation, Virginia Tech).

6. Broxton, C. N., & Culotta, V. C. (2016). SOD enzymes and microbial pathogens: surviving the oxidative storm of infection. PLoS pathogens, 12(1), e1005295.

7. Byrne, B., & Swanson, M. S. (1998). Expression of Legionella pneumophila virulence traits in response to growth conditions. Infection and immunity, 66(7), 3029–3034.

8. Cunha, B. A., Burillo, A., & Bouza, E. (2016). Legionnaires’ disease. The Lancet, 387(10016), 376–385.

9. de Felipe, K. S., Glover, R. T., Charpentier, X., Anderson, O. R., Reyes, M., Pericone, C. D., & Shuman, H. A. (2008). Legionella eukaryotic-like type IV substrates interfere with organelle trafficking. PLoS pathogens, 4(8), e1000117.

10. Demirjian, A., Lucas, C. E., Garrison, L. E., Kozak-Muiznieks, N. A., States, S., Brown, E. W., … & Hicks, L. A. (2015). The importance of clinical surveillance in detecting Legionnaires’ disease outbreaks: A large outbreak in a hospital with a Legionella disinfection system—Pennsylvania, 2011–2012. Clinical Infectious Diseases, 60(11), 1596–1602.

11. Escoll, P., Rolando, M., Gomez-Valero, L., & Buchrieser, C. (2014). From amoeba to macrophages: exploring the molecular mechanisms of Legionella pneumophila infection in both hosts. Molecular mechanisms in Legionella pathogenesis, 1–34.

12. Feeley, J. C., Gibson, R. J., Gorman, G. W., Langford, N. C., Rasheed, J. K., Mackel, D. C., & Baine, W. B. (1979). Charcoal-yeast extract agar: primary isolation medium for Legionella pneumophila. Journal of clinical microbiology, 10(4), 437–441.

13. Fields, B. S., Shotts Jr, E. B., Feeley, J. C., Gorman, G. W., & Martin, W. T. (1984). Proliferation of Legionella pneumophila as an intracellular parasite of the ciliated protozoan Tetrahymena pyriformis. Applied and environmental microbiology, 47(3), 467–471.

14. Gal-Mor, O., & Segal, G. (2003). The Legionella pneumophila GacA homolog (LetA) is involved in the regulation of icm virulence genes and is required for intracellular multiplication in Acanthamoeba castellanii. Microbial pathogenesis, 34(4), 187–194.

15. Giachino, A., & Waldron, K. J. (2020). Copper tolerance in bacteria requires the activation of multiple accessory pathways. Molecular microbiology, 114(3), 377–390

16. Gillmaier, N., Schunder, E., Kutzner, E., Tlapák, H., Rydzewski, K., Herrmann, V., … & Heuner, K. (2016). Growth-related metabolism of the carbon storage poly-3-hydroxybutyrate in Legionella pneumophila. Journal of Biological Chemistry, 291(12), 6471–6482.

17. Government of Canada, P. H. A. of C. (2021, July 20). Reported cases from 1924 to 2019 in Canada - notifiable diseases on-line. Government of Canada, Public Health Agency of Canada. Retrieved October 30, 2022, from https://diseases.canada.ca/notifiable/charts?c=pl

18. Hammer, B. K., Tateda, E. S., & Swanson, M. S. (2002). A two-component regulator induces the transmission phenotype of stationary-phase Legionella pneumophila. Molecular microbiology, 44(1), 107–118.

19. Hammer, B. K., Tateda, E. S., & Swanson, M. S. (2002). A two-component regulator induces the transmission phenotype of stationary-phase Legionella pneumophila. Molecular microbiology, 44(1), 107–118.

20. Hovel-Miner, G., Pampou, S., Faucher, S. P., Clarke, M., Morozova, I., Morozov, P., … & Kalachikov, S. (2009). σS controls multiple pathways associated with intracellular multiplication of Legionella pneumophila. Journal of Bacteriology, 191(8), 2461–2473.

21. Huston, W. M., Naylor, J., Cianciotto, N. P., Jennings, M. P., & McEwan, A. G. (2008). Functional analysis of the multi-copper oxidase from Legionella pneumophila. Microbes and infection, 10(5), 497–503.

22. James, B. W., Mauchline, W. S., Dennis, P. J., Keevil, C. W., & Wait, R. (1999). Poly-3-hydroxybutyrate in Legionella pneumophila, an energy source for survival in low-nutrient environments. Applied and environmental microbiology, 65(2), 822–827.

23. Khan, S., Beattie, T.K. & Knapp, C.W. Rapid selection of antimicrobial-resistant bacteria in complex water systems by chlorine and pipe materials. Environ Chem Lett 17, 1367–1373 (2019). 10.1007/s10311-019-00867-z

24. Kim, E. H., Charpentier, X., Torres-Urquidy, O., McEvoy, M. M., & Rensing, C. (2009). The metal efflux island of Legionella pneumophila is not required for survival in macrophages and amoebas. FEMS microbiology letters, 301(2), 164–170.

25. Kim, E. H., Charpentier, X., Torres-Urquidy, O., McEvoy, M. M., & Rensing, C. (2009). The metal efflux island of Legionella pneumophila is not required for survival in macrophages and amoebas. FEMS microbiology letters, 301(2), 164–170.

26. Kunz, J. M. (2024). Surveillance of Waterborne Disease Outbreaks Associated with Drinking Water—United States, 2015–2020. MMWR. Surveillance Summaries, 73.

27. Lawrence A, K Nicholls S, H Stansfield S, M Huston W. 2014. Characterization of the tail-specific protease (Tsp) from Legionella. J Gen Appl Microbiol 60:95–100.

28. LeChevallier, M. (2023). Examining the efficacy of copper-silver ionization for management of Legionella: Recommendations for optimal use. AWWA Water Science, 5(2), e1327.

29. Li, L., Mendis, N., Trigui, H., & Faucher, S. P. (2015). Transcriptomic changes of Legionella pneumophila in water. BMC genomics, 16, 1–21.

30. Liang, J., & Faucher, S. P. (2024). Interactions between chaperone and energy storage networks during the evolution of Legionella pneumophila under heat shock. PeerJ, 12, e17197.

31. Liang, J., Cameron, G., & Faucher, S. P. (2023). Development of heat-shock resistance in Legionella pneumophila modeled by experimental evolution. Applied and Environmental Microbiology, 89(9), e00666–23.

32. Lin, Y. S. E., Vidic, R. D., Stout, J. E., & Victor, L. Y. (1996). Individual and combined effects of copper and silver ions on inactivation of Legionella pneumophila. Water Research, 30(8), 1905–1913.

33. Liu, Z., Stout, J. E., Tedesco, L., Boldin, M., Hwang, C., Diven, W. F., & Yu, V. L. (1994). Controlled evaluation of copper-silver ionization in eradicating Legionella pneumophila from a hospital water distribution system. Journal of Infectious Diseases, 169(4), 919–922.

34. Lynch, D., Fieser, N., Glöggler, K., Forsbach-Birk, V., & Marre, R. (2003). The response regulator LetA regulates the stationary-phase stress response in Legionella pneumophila and is required for efficient infection of Acanthamoeba castellanii. FEMS microbiology letters, 219(2), 241–248.

35. Lynn, D. H., & Doerder, F. P. (2012). The life and times of Tetrahymena. Methods in cell biology, 109, 9–27.

36. Marston, B. J., Lipman, H. B., & Breiman, R. F. (1994). Surveillance for Legionnaires’ disease: risk factors for morbidity and mortality. Archives of Internal Medicine, 154(21), 2417–2422.

37. Matthews, S., Trigui, H., Grimard-Conea, M., Vallarino Reyes, E., Villiard, G., Charron, D., … & Prevost, M. (2022). Detection of diverse sequence types of Legionella pneumophila by legiolert enzymatic-based assay and the development of a long-term storage protocol. Microbiology Spectrum, 10(6), e02118–22.

38. McMullen, C. K., Dougherty, B., Medeiros, D. T., Yasvinski, G., Sharma, D., & Thomas, M. K. (2024). Estimating the burden of illness caused by domestic waterborne Legionnaires’ disease in Canada: 2015–2019. Epidemiology & Infection, 152, e18.

39. Molofsky AB, Swanson MS. Legionella pneumophila CsrA is a pivotal repressor of transmission traits and activator of replication. Mol Microbiol. 2003 Oct;50(2):445–61. doi: 10.1046/j.1365-2958.2003.03706.x. PMID: 14617170.

40. Molofsky, A. B., & Swanson, M. S. (2004). Differentiate to thrive: lessons from the Legionella pneumophila life cycle. Molecular microbiology, 53(1), 29–40.

41. Morel FMM, Westall JC, Ruerer JG, Chaplick JP. Descriptions of algal growth media AQUIL and FRAQUIL. Technical Note, R M Parsons Laboratory for Water Resources and Hydrodynamics Massachusetts Institute of Technology, Cambridge, MA. 1975.

42. Nanney, D. L., & McCoy, J. W. (1976). Characterization of the species of the Tetrahymena pyriformis complex. Transactions of the American Microscopical Society, 664–682.

43. Nicolau, A., Mota, M., & Lima, N. (1999). Physiological responses of Tetrahymena pyriformis to copper, zinc, cycloheximide and Triton X-100. FEMS microbiology ecology, 30(3), 209–216.

44. Rasis, M., & Segal, G. (2009). The LetA-RsmYZ-CsrA regulatory cascade, together with RpoS and PmrA, post-transcriptionally regulates stationary phase activation of Legionella pneumophila Icm/Dot effectors. Molecular microbiology, 72(4), 995–1010.

45. Rowbotham, T. J. (1980). Preliminary report on the pathogenicity of Legionella pneumophila for freshwater and soil amoebae. Journal of clinical pathology, 33(12), 1179–1183.

46. Rowbotham, T. J. (1980). Preliminary report on the pathogenicity of Legionella pneumophila for freshwater and soil amoebae. Journal of clinical pathology, 33(12), 1179–1183.

47. Rowland, J. L., & Niederweis, M. (2013). A multicopper oxidase is required for copper resistance in Mycobacterium tuberculosis. Journal of bacteriology, 195(16), 3724–3733.

48. Sahr, T., Rusniok, C., Impens, F., Oliva, G., Sismeiro, O., Coppee, J. Y., & Buchrieser, C. (2017). The Legionella pneumophila genome evolved to accommodate multiple regulatory mechanisms controlled by the CsrA-system. PLoS genetics, 13(2), e1006629.

49. Salazar-Ardiles, C., Asserella-Rebollo, L., & Andrade, D. C. (2022). Free-living amoebas in extreme environments: the true survival in our planet. BioMed Research International, 2022(1), 2359883.

50. Saoud, J., Mani, T., & Faucher, S. P. (2021). The tail-specific protease is important for Legionella pneumophila to survive thermal stress in water and inside amoebae. Applied and Environmental Microbiology, 87(9), e02975–20.

51. Singh, S. K., Parveen, S., SaiSree, L., & Reddy, M. (2015). Regulated proteolysis of a cross-link–specific peptidoglycan hydrolase contributes to bacterial morphogenesis. Proceedings of the National Academy of Sciences, 112(35), 10956–10961.

52. Solioz, M. (2018). Copper and bacteria: evolution, homeostasis and toxicity (p. 88). Cham, Switzerland: Springer International Publishing.

53. Storey, M. V., Winiecka-Krusnell, J., Ashbolt, N. J., & Stenström, T. A. (2004). The efficacy of heat and chlorine treatment against thermotolerant Acanthamoebae and Legionellae. Scandinavian journal of infectious diseases, 36(9), 656–662.

54. Tanner, J. R., Li, L., Faucher, S. P., & Brassinga, A. K. C. (2016). The CpxRA two-component system contributes to L egionella pneumophila virulence. Molecular Microbiology, 100(6), 1017–1038.

55. Trainer MA, Charles TC. The role of PHB metabolism in the symbiosis of rhizobia with legumes. Appl Microbiol Biotechnol. 2006;71(4):377–86.

56. Trigui, H., Dudyk, P., Sum, J., Shuman, H. A., & Faucher, S. P. (2013). Analysis of the transcriptome of Legionella pneumophila hfq mutant reveals a new mobile genetic element. Microbiology, 159(Pt_8), 1649–1660.

57. van Loosdrecht MCM, Pot MA, Heijnen JJ. Importance of bacterial storage polymers in bioprocesses. Water Sci Technol. 1997;35(1):41–7.

58. Xu, F., Liu, S., Naren, N., Li, L., Ma, L. Z., & Zhang, X. X. (2022). Experimental evolution of bacterial survival on metallic copper. Ecology and Evolution, 12(8), e9225.

59. Yamamoto, K., & Ishihama, A. (2006). Characterization of copper-inducible promoters regulated by CpxA/CpxR in Escherichia coli. Bioscience, biotechnology, and biochemistry, 70(7), 1688–1695.

60. Yu, A. T., Kamali, A., & Vugia, D. J. (2019). Legionella epidemiologic and environmental risks. Current Epidemiology Reports, 6, 310–320.

