## Supplementary Figures for "Copper resistance in *Legionella pneumophila*: role of genetic factors and host cells"

### <u>Supplemental Figures for Copper resistance in *Legionella pneumophila*: role of genetic factors and host cells.</u>

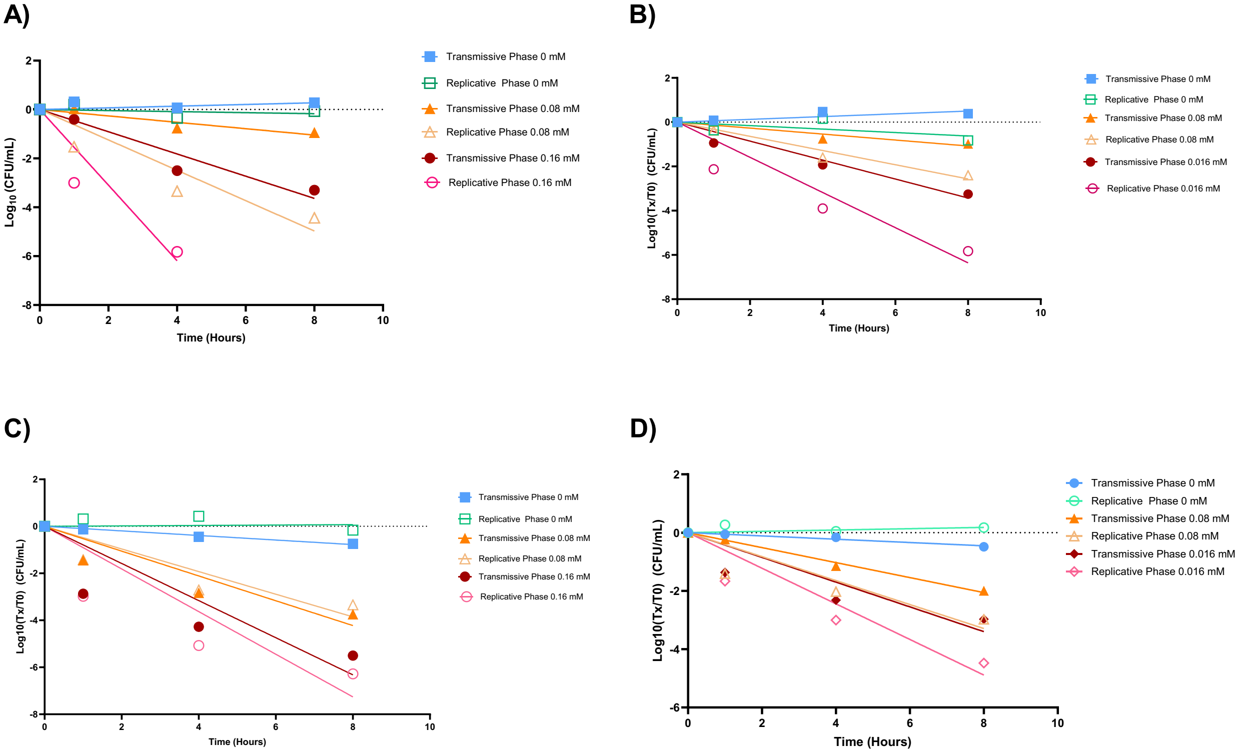


**Supplementary Figure 1:** Individual biological replicates of reduction in cell concentration over time of *L. pneumophila* Philadelphia-1 in transmissive phase and replicative phase. PE cultures were grown at 37°C for 48 hours, while E phase cultures were grown for 12 hours at 37°C. Cultures were washed, resuspended, and diluted in Fraquil to a final concentration of $10^8$ CFU/mL. The bacterial suspensions were incubated overnight at room temperature, then exposed to 0, 0.08, and 0.16 mM $CuCl_2$. CFU counts were taken at 0, 1, 4, and 8 hours. The CFU count data was baseline corrected, transformed, and fitted with a line of best fit. Each biological replicate was done in triplicate.

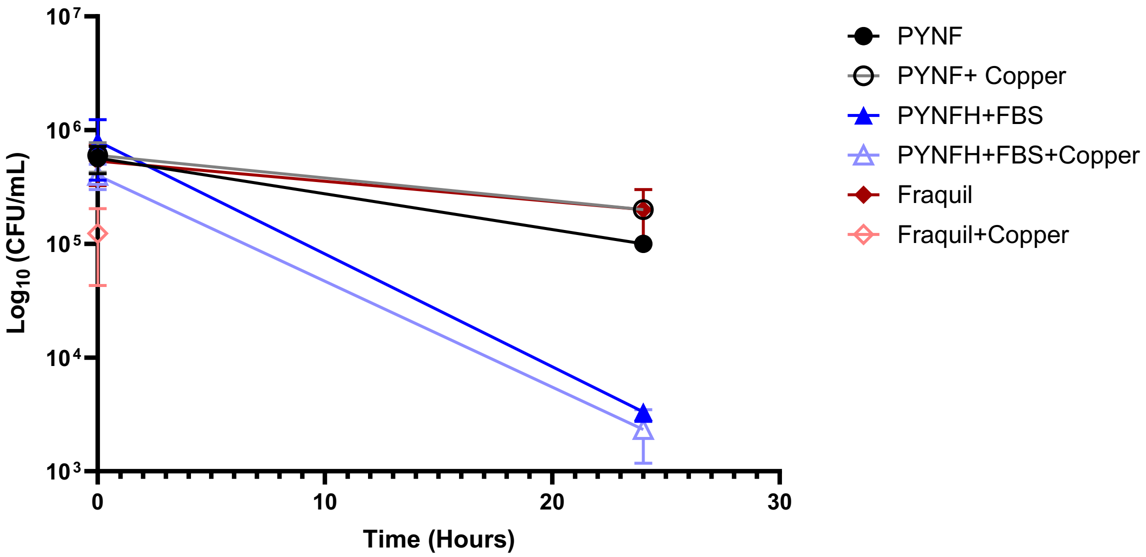


**Supplementary Figure 2:** Comparison of *L. pneumophila* Philadelphia-1 survival in coculture media with and without copper. A final concentration of $10^5$ cells/mL *L. pneumophila* Philadelphia-1 was added to a 24 well plate. For the copper exposed conditions, a final concentration of 0.08 mM $CuCl_2$ was added to each well. 985 uL of media was then added to the wells. The plate for the 24-hour conditions were incubated at 37ºC for 24 hours.

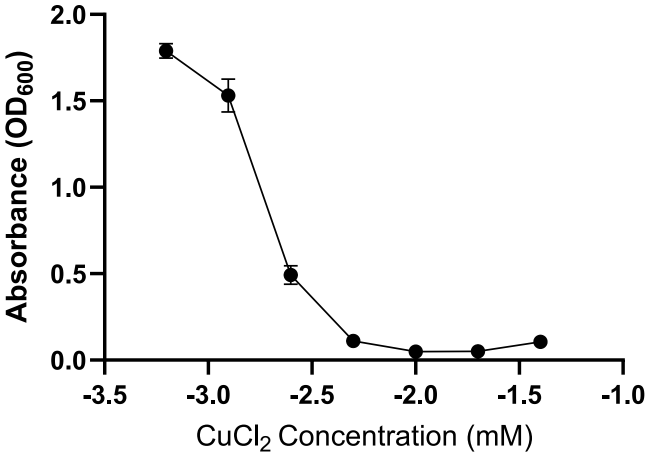


**Supplementary Figure 3:** Minimum inhibitory concentration (MIC) of $CuCl_2$ for *L. pneumophila* Philadelphia-1. $CuCl_2$ was serially diluted in a 96 well plate in AYE media. No copper was added to the last row of the 96 well plate. 20 µL of a Philadelphia-1 suspension with a concentration of $10^8$ cells/mL was added to each well. The 96 well plate was incubated for 72 hours at 37°C. Absorbance was measured to compare cell growth at each concentration. At $CuCl_2$ concentrations higher than 2 mM, the copper ions were detected at $OD_{600}$ despite no visible growth in the corresponding well. Absorbance data transformed with X=log (x)
